# Two major-effect loci influence assortative mating in females of the sibling species, *Drosophila simulans* and *D. sechellia*

**DOI:** 10.1101/2024.04.30.591936

**Authors:** Kenneth Lu, Deniz Erezyilmaz

## Abstract

Secondary contact between incompletely isolated species can produce a wide variety of outcomes, including production of new species and adaptive radiations. The vinegar flies *Drosophila simulans* and *D. sechellia* diverged on islands in the Indian Ocean and are currently separated by partial pre– and postzygotic barriers. The recent discovery of hybridization between *D. simulans* and *D. sechellia* in the wild presents an opportunity to monitor the prevalence of alleles that influence introgression between these species. We therefore sought to identify those loci that affect assortative mating, and we adapted a two-choice assay to test behavioral isolation in females. Using high-resolution seq-based QTL mapping, we found two major effect loci on the third chromosome that have a profound effect upon assortative mating of females. Each major QTL accounts for 32-37% of the difference in phenotype on its own, which is highly significant for a behavioral trait. The two major QTL of both backcrosses co-localize in one-dimensional analyses, suggesting that they may be alternate alleles of the same loci. The major-effect loci also co-localize with genes that encode enzymes required for female– and species-specific production of the pheromone, 7,11-heptacosadiene, emphasizing the importance of female attractiveness to males in separation of these species. Moreover, the genetic architecture of female assortative mating may be a factor in species separation, since alleles that influence assortative mating in females are linked to major-effect loci that influence assortative mating in males, and to loci that contribute to host fruit adaptation in *D. sechellia*.

## Introduction

Closely related species that have been evolving independently often retain the ability to interbreed, despite some degree of pre– or post-zygotic isolation. When such divergent species come back into contact they may become more distinct through ‘reinforcement’ (Coyne and Orr 1989; Coyne and Orr 1997), blend together through ‘de-speciation’ (Turner 2002), form an entirely new lineage through ‘hybrid speciation’ (Lamichhaney et al. 2018; Mallet 2007), or they may persist as long-term stable ‘hybrid swarms’ (Barton and Hewitt 1989; Pfennig 2007; Servedio and Hermisson 2020). Which of these distinct outcomes will transpire may depend, in large part, upon the sensory cues that related species use to recognize conspecific mates, and upon their genetic basis.

The nascent species, *Drosophila simulans* and *D. sechellia* are distinguished by host plant specialization and assortative mating. *D. simulans* is a cosmopolitan generalist that probably first emerged in Madagascar and utilizes a wide variety of rotting fruit. *D. sechellia* is endemic to the Seychelles islands where it eats and breeds primarily on the fruit of *Morinda citrifolia*, which is toxic to all other Drosophila, including *D. simulans* (R’Kha et al. 1991). In laboratory tests the two species will interbreed, but behavioral barriers affect males and females differently. *D. simulans* females and *D. sechellia* males readily mate in no-choice assays, but the reciprocal cross is rarely successful (Lachaise et al. 1986). Speciation between *D. simulans* and *D. sechellia* has progressed to partial postzygotic isolation, since F1 male offspring are sterile although F1 females are fertile (Coyne et al. 1991). The two species are currently interbreeding in the Seychelles and wild-caught hybrid flies have Y chromosomes from *D. sechellia* and mitochondria from *D. simulans*, indicating that the same mating asymmetry is also a factor in wild populations (Matute and Ayroles 2014).

Chemosensory cues conferred by distinct non-volatile contact pheromones produced by specialized cells in the abdomens of females of each species are necessary for the barrier between *D. simulans* males and *D. sechellia* females (Billeter et al. 2009; Coyne et al. 1994a; Coyne and Oyama 1995). An unsaturated 23 carbon with a double bond at C7, 7-Tricosene (7-T), is the predominant contact pheromone produced by *D. simulans* males and females, while *D. sechellia* females produce the 27-carbon compound, 7,11-heptacosadiene with a second double bond on C11 (7,11-HD; Jallon and David 1987). 7,11-HD on females is detected by contact chemoreceptors on the male forelimb (Thistle et al. 2012). *D. simulans* males do not court *D. sechellia* females after contact, but when the forelimb of *D*. *simulans* males is removed they will sustain courtship towards *D. sechellia* females (Shahandeh et al. 2018). *D. melanogaster* females also produce 7,11-HD like *D. sechellia* females. However, crosses between *D. simulans* and *D. melanogaster* occur more readily when the female is *D. melanogaster* and the male is *D. simulans* than with the reciprocal cross (Carracedo et al. 1998), although this trait can vary among strains (Izquierdo et al. 1992; Gérard and Presgraves 2012). In any case, if *D. simulans* males will mate with *D. melanogaster* females but not *D. sechellia* females, this shows that 7,11-HD production in *D. sechellia* females in not the only barrier that separates *D. simulans* males from *D. sechellia* females.

Auditory cues also contribute to isolation between *D. simulans* and *D. sechellia*. Male fruit flies from the *D. simulans* complex ‘sing’ species-specific courtship songs by vibrating their wings (Gleason and Ritchie 2004; Ritchie et al. 1999). *D. simulans* females mate more quickly when stimulated by the song of *D. simulans males* than *D. sechellia* males (Ritchie et al. 1999), and *D. sechellia* females are more likely to mate with *D. simulans* males if their wings are removed; apparently no song is better than producing the wrong song in inducing female receptivity (Tomaru et al. 2004). Assortative mating in this species pair therefore occurs through a combination of male preference and female choice, conferred through chemosensory and auditory cues, respectively.

Given the importance of this genetic model of hybridizing species, we have conducted this current study using high-resolution seq-based genetic mapping to reveal those loci that prevent or facilitate interbreeding. We established a two-choice assay that uses copulation to test isolation between *D. simulans* and *D. sechellia* females from heterospecific males. Assortative mating in females is comprised of female preferences and those factors that are used as a basis of preference, and both these traits will be captured in our assay. We find that the genetic basis of assortative mating in females is localized to two QTL on different arms of the third chromosome, linked by epistasis and co-localizing with loci that encode enzymes required for production of 7,11-HD. The effects of the two major QTL confer aspects of isolation and hybridization. For instance, *D. simulans* alleles at QTL-3L and QTl-3R increased the likelihood of copulation with *D. simulans* males, but females that were hybrid at both the major QTLs were more likely to mate with *D. sechellia* males when given a choice. Finally, we suggest that the proximity of the female assortative mating loci to loci that confer assortative mating in males on chromosome 3R as well as loci that confer resistance to *M. citrifolia* toxin could have been a factor in the separation of *D. simulans* and *D. sechellia* as well as their ongoing hybridization in the Seychelles.

## Materials & Methods

### Drosophila strains

The *D. sechellia^13^* strain (Tucson Stock Center #14021-0248.13) is derived from flies captured on Cousin Island in the Seychelles, an undeveloped nature preserve where *D. sechellia* was first discovered. *D. sechellia^D1A1C^* is an inbred strain that was produced by sib-mating single females and males of the *D. sechellia^13^* strain for five generations. The *D. simulans* Tsimbazaza strain, which we obtained from David Stern (Janelia, HHMI), was originally isolated from Madagascar, where *D. simulans* variation is greatest and is probably where the species originated (Kopp et al. 2006). The *D. simulans^A2A2B^*was produced by sib-mating single females and males for five generations.

### Mate choice assays

For the two-choice tests, virgin adult flies were sexed at eclosion and cultured together in groups of three individuals of the same sex in vials without yeast. We used females that were older than 3 days but less than 28 days old, and males that were between 7 and 28 days old in mate choice tests. For the mating assays, live yeast paste was added to empty fly food vials (Genesee Scientific, San Diego, CA). Flies were cultured at 25C on a 12:12 light:dark cycle and the assays commenced within one hour of subjective dawn. The assay began when three virgin females were combined with three *D. simulans* males and three *D. sechellia* males. All six of the males in each test vial were aged the same number of days since eclosion. The assay was monitored for four hours or until copulation, whichever came first. Once the first pair mated, the vial was moved to the freezer (–20C), so that the pair were frozen *in copulo*. The species of copulating male was determined by examining the male genitalia, which is distinct for *D. simulans* and *D. sechellia* (Liu et al. 1996; Lu 2022). To test if female age were a factor in interspecific mating of *D. simulans^A2A2B^* females, we set up eleven trials of 23-24 days old *D. simulans^A2A2B^*. Females in five of six *D. simulans^A2A2B^* trials mated with *D. simulans^A2A2B^* males, which is not different from the ratio observed in our larger study and it is consistent with the interspecific mating observed by others.

For the no-choice tests, males and females were isolated and cultured in the same way as for the two choice tests, but three female progeny from backcrosses were combined with males from just one species. For these tests, we ended the trial after two hours, rather than four.

### Production of backcross populations

F1 females were first produced by crossing a *D. simulans^A2A2B^* female to a *D. sechellia^13^* male, because the reciprocal cross produces few offspring. F1 females were backcrossed to either *D. simulans^A2A2B^*males to produce *D. simulans* backcross progeny, or to *D. sechellia^13^* males to produce *D. sechellia* backcross progeny.

### Production of Multiplexed Shotgun Genotyping Libraries

We used multiplexed shotgun genotyping (MSG) to infer the ancestry of individual female backcross progeny at hundreds of thousands of loci (Andolfatto et al. 2011; Cande et al. 2012). As described previously (Andolfatto et al. 2011), genomic DNA was extracted from individual flies in 96 well plates using the Puregene Tissue Kit (Qiagen, Venlo, Netherlands). Each Illumina library was comprised 384 individual barcoded genomes. Single-end 100bp sequencing was performed on an Illumina Hi-Seq at the University of Oregon Genomics and Cell Characterization Facility (Eugene, Oregon). To create a more customized reference genome we separately updated the *D. simulans* genome assembly (Hu et al. 2013) with paired-end sequence from specific strains, *D. sechellia^D1A1C^* and *D. simulans^A2A2B^* using the script, msgUpdateParental.pl, which is a part of the msg package (Andolfatto et al. 2011). The paired-end libraries of *D. sechellia^D1A1C^* and *D. simulans^A2A2B^* gDNA were created at U. of Oregon G3 genomics Center and run on an Illumina Hi-Seq 2000 sequencer.

### Genotyping and QTL Mapping

The MSG package was installed on the cluster at Janelia Farm. We first ran each of four libraries of 384 individuals and individuals with poor quality genotyping were eliminated from the final analysis. We then combined the parsed individual files into one large dataset and the entire set for each backcross was re-run together. The file of *D. simulans* backcross female progeny contained 782 individual genomes and was genotyped at 654,693,365 markers (Lu 2022b). We used the script pull_thin (Stern 2018; Cande et al. 2012), which thins the dataset to only include markers that flank a recombination breakpoint in at least one individual, to thin the *D. simulans* backcross genotype file to 3,461 markers. For the *D. sechellia* backcross the file of 690 individual female genomes was genotyped at 547,992,323 markers, then similarly thinned to 2,890 markers (Lu 2022a). The phenotype was treated as a binary trait; those females that mated with males of the same species as males of the back-cross (their father’s species) were scored ‘0’ and those females that mated with males from the second species were scored with a ‘1’.

We used the R/qtl package (Broman et al. 2003) to analyze our backcross libraries. The function, *scanone* was used for one-dimensional analysis, and *cim* was used for composite interval mapping. The binary model was used for each analysis. We used the Haley-Knott algorithm for interval mapping, composite interval mapping, and model building. 1000 permutation replicates were used to assess the statistical significance of our log of the odds (LOD) scores for both the *scanone* and *cim* functions. Significance intervals were inferred from a 2-LOD score drop from the peak LOD value. *fitqtl* and *stepwiseqtl* were used to explore multiple QTL models and pairs of interactions using forward selection and backcross elimination (Broman et al. 2003).

For PCR-based genotyping, we used the primer pairs *desatF_for/desatF_rev* (Tm = 60) and *eloF_for/eloF_rev* (Tm = 62) with MyTaq Red (Bioline, Meridian Bioscience) to distinguish between *D. simulans* and *D. sechellia* alleles. The primer sequences *eloF* 5’ CAACATATTCCAGATCCTTTACAA3’ e*loF*-R 5’ ATCCTTATATTTGTGATCCATCG3’, which amplify a species-specific sequence length polymorphism at ∼3R:15.39 Mbp. The primers *desatF*-F CCTGAACACTTTGGCCTTCC and *desatF*-R 5’ATTTGCTTGCCCTTCTCCAC3’ (Swartzlander 2016), amplify a species-specific sequence length polymorphism at ∼3L:10.76 Mbp. We used Prism software to assess the significance of associations in 2×2 Contingency Tables using Fisher’s Exact Test genotype frequencies between mating phenotypes.

To measure epistasis, we tested the fit of the assortative mating data from our *D. simulans* backcross mapping population to an additive model, (a + d) – (b + c) = 0, where a = the proportion of females that mated with D*. sechellia* males with the e/i;e/i genotype, b = the proportion of females that mated with *D. sechellia* males with the e/i;i/I genotype, c = the proportion of females that mated with *D. sechellia* males with the i/i;e/i genotype and d = the proportion of females that mated with *D. sechellia* males with the i/i;i/i genotype. We approximated the standard error assuming a Gaussian distribution of outcomes. The complete calculation is given in supplementary fileS1.

## Results

We devised a high throughput method to screen for alleles that contribute to the behavioral barrier between *D. simulans* and *D. sechellia* (Figure 1A). We used a two-choice assay, a standard test of behavioral isolation (Bay et al. 2017; Boake et al. 2003; Yukilevich and True 2008), instead of a no-choice assay because inter-specific mating occurs more readily than otherwise when flies are not given a choice (Coyne et al. 2005). This approach has the advantage that each “choice” represents a positive unambiguous score of behavioral isolation. Copulation was too infrequent when females were tested one-by-one, so three females were combined with three *D. simulans* males and three *D. sechellia* males for each mating trial, and these were observed for four hours. We included live yeast paste in the test vial because it stimulates female receptivity (Gorter et al. 2016). This simple two-choice design captures both female preference loci and loci that confer female attractiveness to males.

**Figure 1.**
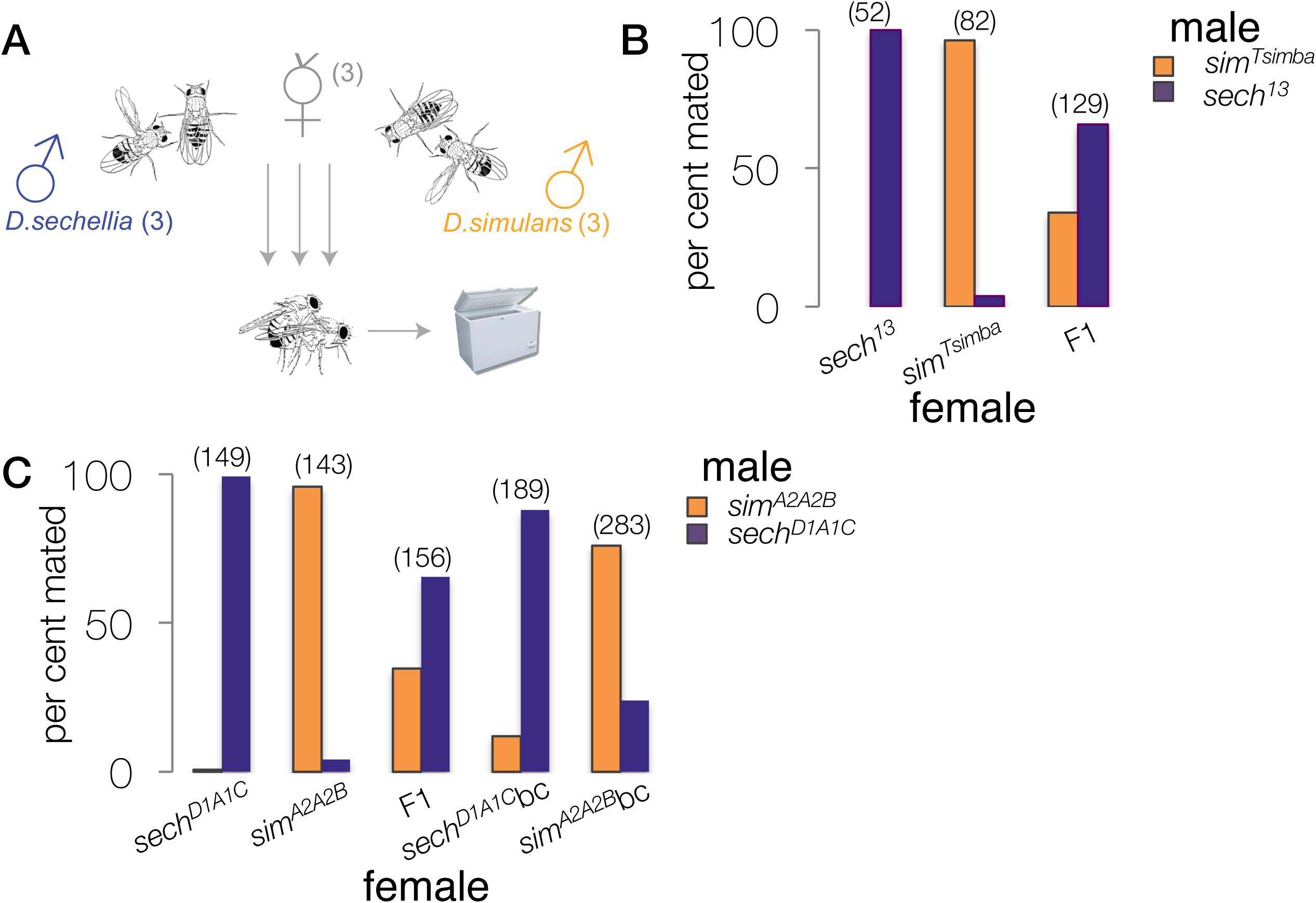
**A**. Design of the two-choice test of assortative mating in females. Each vial contained: 1) three test females, (either pure species, F1 females or F1 backcross progeny), 2) three *D. sechellia* males and 3) three *D. simulans* males. When the first pair copulates, the vial is moved to a freezer at –20C. Once the copulating pair is frozen stiff, the species of copulating male is determined by examining the male genitalia, which is distinct for each species. **B.** Assortative mating of female from the Tsimbazaza strain of *D. simulans* (*sim^Tsimba^*) and females of *D. sechellia* strain (*sech^13^*). **C**. Assortative mating of females from inbred lines derived from *D. simulans^Tsimba^* and *D. sechellia^13^*, the F1 progeny and the F1 back-cross progeny. The number of copulating pairs is given in parentheses.

We examined the strength of assortative mating in females of the pure species, inbred lines, hybrids and backcross progeny (Figure 1B). Copulation between a *D. sechellia^1^*^3^ female and *D. simulans^Tsimba^*male was very rare in the two-choice test, but *D. sechellia* males and *D. simulans* females were first to copulate in approximately 5% of the trials. We learned that F1 hybrid female progeny, from a *D. simulans^Tsimba^* mother and *D. sechellia^13^* father, mate more frequently with *D. sechellia^13^* males by nearly a 2:1 ratio indicating partial dominance for this trait (Figure 1B). These results resemble what has been previously reported with other strains (Lachaise et al. 1986; Cobb and Jallon 1990; Coyne 1992). We then confirmed that these phenotypes hold in our mapping strains, *D. sechellia^D1A1C^* and *D. simulans^A2A2B^*, inbred lines of *D. simulans^Tsimba^*and *D. sechellia^13^* strains respectively. While 0 of 149 *D. sechellia^D1A1C^* females mated with *D. simulans^A2A2B^* males, some *D. simulans^A2A2B^*females mated with *D. sechellia^D1A1C^* males, reflecting the pattern that was observed in the pure species (Figure 1C). We backcrossed F1 females to produce 189 *D. sechellia* backcross progeny and separately, 283 female progeny of the *D. simulans* backcross. These females tended to favor males that were the same species as their father (Figure 1C).

### QTL mapping of assortative mating in females of the *D. simulans* backcross

A second cohort of female progeny of the *D. simulans* backcross were collected and tested in the two-choice assay. In this case, the three females were heterogeneous recombinants, which added competition among the female genotypes to the assay, selecting for the strongest association with *D. simulans* or *D. sechellia* males. Once roughly equal numbers of each phenotype (mating with either species of male) were tested, we processed individual genomes for MSG analysis (Andolfatto et al. 2011). MSG produced genotypes at hundreds of thousands of markers, which we then thinned to only those markers that straddle a recombination breakpoint in at least one individual (Cande et al. 2012; Stern 2018). We treated copulation as a binary trait and mapped 415 recombinant backcross females that mated with *D. simulans* males and 367 genomes of recombinant backcross females that had mated with *D. sechellia* males. This analysis reveals two highly significant QTL peaks: one centered at ∼3L:11.31 Mbp with LOD = 19.86 (Figure 2A), and another, larger peak (LOD = 25.08) is centered over a marker near ∼ 3R:14.75 Mbp (Figure 2A). The entire third chromosome was significant, so we next performed composite interval mapping to reduce residual variation (grey line, Figure 2A). This analysis suggested that in addition to the main QTL on 3R, an additional significant locus is found near the distal end of the chromosome, ∼3R:22.15 Mbp (LOD = 3.07).

**Figure 2.**
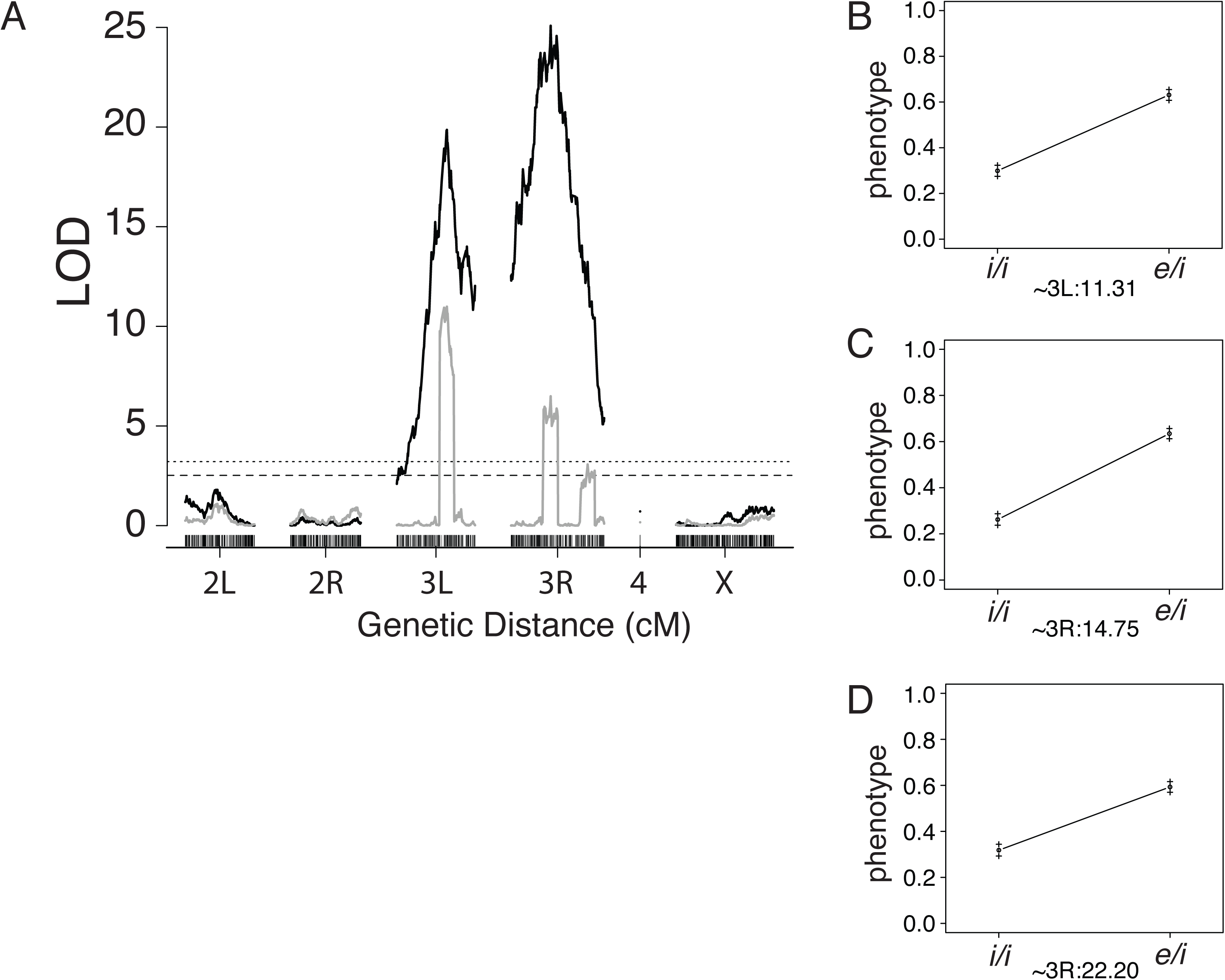
A. Results of one-dimensional analysis of interval mapping (black lines) and composite interval mapping (grey lines) for. *D. simulans* backcross. Chromosome markers (are given on the x-axis over genetic distance with log of the odds ratio (LOD) on the y-axis. The horizontal dashed lines indicate the *p* = 0.05 significance threshold, and the dotted line shows the *p =* .01 significance threshold. **B-D.** Effect plots showing the phenotypic effect of substituting a *D. sechellia* allele (*e*) for a *D. simulans* allele (*i*) where phenotype is 0 = mating with a *D. simulans* male and 1 = mating with a *D. sechellia* male at the markers ∼3L:11.31 (**B**) ∼3R:14.75 (**C**) and ∼3R:22.20 (**D**). The standard error is shown in the error bars.

Because our analyses indicated that more than one locus contributes to assortative mating in females, we next examined the fit of multiple QTL models (Table 1). We used forward selection and backwards elimination of all models containing between two to five QTL and searched for interactions between each pair of QTL. The best-fit model includes the three major-effect loci that were identified in our one-dimensional analysis: ∼3L:11.31 Mbp (QTL-3L*_sim_*) and the two loci on the right arm of chromosome three, ∼3R:14.76 Mbp (QTL-3R*_sim_*A) and ∼3R:22.15 (QTL-3R*_sim_*B; Figure 2A, Table 1). The model also includes a significant interaction between QTL-3L*_sim_* and QTL-3R*_sim_*B. The effect of either locus on their own is considerable; substitution of a *D. sechellia* allele at the locus on 3L results in 32% increase in the likelihood of mating with a *D. sechellia* male and substitution of a *D. sechellia* allele at QTL-3R*_sim_*A increases the likelihood of mating with *D. sechellia* males by 37%. At QTL-3R*_sim_*B, substitution of a *D. sechellia* allele has an effect of 22.3% (Figure 2B, C, D).

**Table 1.** QTL locations and best-fit models for QTL analyses of the *D. simulans* backcross (*bc-sim*) and the *D. sechellia* backcross (*bc-sech*).

| Cross | # QTL | % variance (model) | LOD of Model | QTL | QTL Location (Mbp) (2-LOD interval)* | LOD Drop One† | % variance |
| --- | --- | --- | --- | --- | --- | --- | --- |
| *bc-sim* | 3 | 21.6 | 40.30 | QTL-3L<sub>sim</sub> | ~ 3L:11.31<br>(~3L: 10.66 -11.92) | 13.23 | 6.55 |
|  |  |  |  | QTL-3R<sub>simA</sub> | ~ 3R:14.76<br>(~ 3R:14.14 -15.06) | 6.75 | 3.28 |
|  |  |  |  | QTL-3R<sub>simB</sub> | ~ 3R:22.15<br>(~ 3R:21.29 -22.99) | 5.32 | 2.51 |
|  |  |  |  | QTL-3L<sub>sim</sub> *<br>QTL-3R<sub>simB</sub> | - | 2.05 | 1.05 |
| *bc-sech* | 2 | 16.6 | 27.27 | QTL-3L<sub>sech</sub> | ~ 3L: 10.48<br>(~ 3L: 8.53 – 11.90) | 13.96 | 8.14 |
|  |  |  |  | QTL-3R<sub>sechA</sub> | ~ 3R: 16.63<br>(~ 3R:14.55 – 17.43) | 9.82 | 5.65 |

\*The 2-LOD support interval where the LOD score is within 2 LOD of the peak maxima.
†Log-likelihood ratios comparing the full model to a model with the specified QTL removed.

We found significant epistasis between QTL-3L*_sim_* and QTL-3R*_sim_*B in our model building (Table 1). Epistasis between QTL-3L*_sim_* and QTL-3R*_sim_*A, however, was most apparent among those females that had mated with *D. sechellia* males (Figure 4A). Female *D. simulans* backcross progeny that mated with *D. sechellia* were not more likely to have a single *D. sechellia* allele at either QTL but were 3.5 to 5.5 times more likely to bear *D. sechellia* alleles at both QTL-3L*_sim_* and QTL-3R*_sim_*A (Figure 4A). These data are extremely significant, (the effect of one vs two *D. sechellia* alleles at QTL-3L*_sim_* and QTL-3R*_sim_*A was extremely unlikely to be produced by an additive model, *p* = 10^-17^). This pattern shows that for mating with *D. sechellia* males, the effect of a *D. sechellia* allele at only one of the major QTL had no effect on its own; only when both QTL-3L*_sim_* and QTL-3R*_sim_*A bore *D. sechellia* alleles (*e/i; e/i* Figure 4A, right) did we find a shift in the likelihood of mating with *D. sechellia*.

### Alleles that influence assortative mating in females of the *D. sechellia* backcross

We repeated the two-choice assay for a new cohort of female progeny of the *D. sechellia* backcross. DNA of individual genomes were extracted in 96-well plates, sequenced, and processed using the MSG pipeline as we had done for the *D. simulans* backcross. We performed a one-dimensional scan of 287 female progeny of the *D. sechellia* backcross that had mated with *D. simulans* and 403 that had mated with *D. sechellia* males (Figure 1B solid lines). We found that, like the *D. simulans* backcross, the effect of female assortative mating is largely restricted to chromosome three (although a region on chromosome arm 2L was nearly significant at *p* = 0.05, LOD = 2.41), with a strongly significant peak on each arm of the third chromosome. One highly significant locus on 3L at ∼10.75 Mbp (LOD = 17.44), and another on the right arm of the chromosome, ∼3R:16.63 Mbp (LOD = 11.68; Figure 3A).

**Figure 3.**
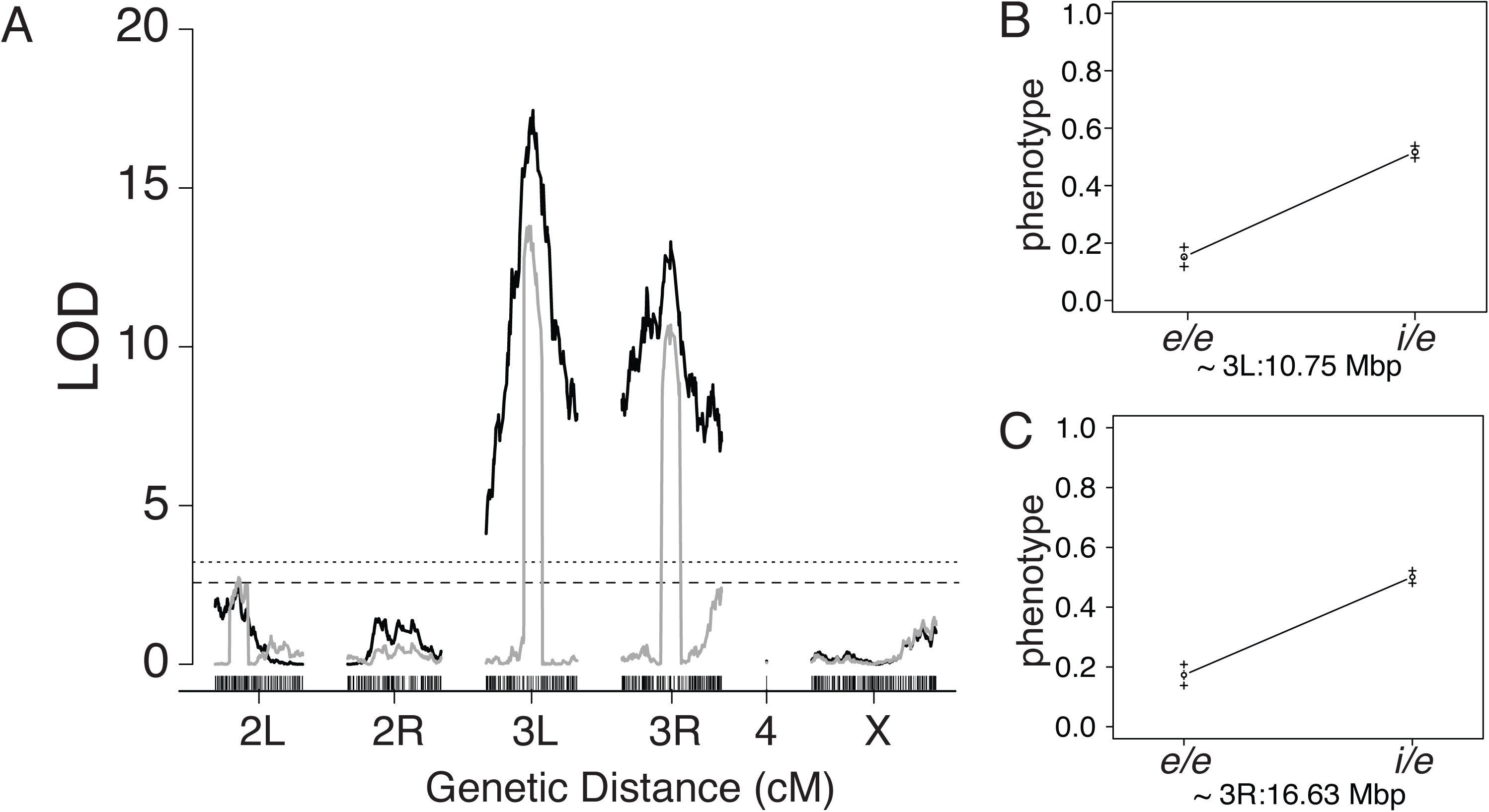
**A**. QTL analysis assortative mating of females among the *D. sechellia* backcross progeny, with interval mapping in black lines and composite interval mapping shown in grey lines. Chromosome markers are given on the x-axis over genetic distance with log of the odds ratio (LOD) on the y-axis. The horizontal dashed lines indicate the *p* = 0.05 significance threshold, and the dotted line shows the *p =* .01 significance threshold. **B, C** Effect plots showing the phenotypic effect of substituting a *D. simulans* allele (*i*) for a *D. sechellia* allele (*e*) where phenotype is 0 = mating with a *D. sechellia* male and 1 = mating with a *D. simulans* male at the markers ∼3L:10.75 (**B**) and ∼3R:16.63 (**C**). The standard error is shown in the error bars.

**Figure 4.**
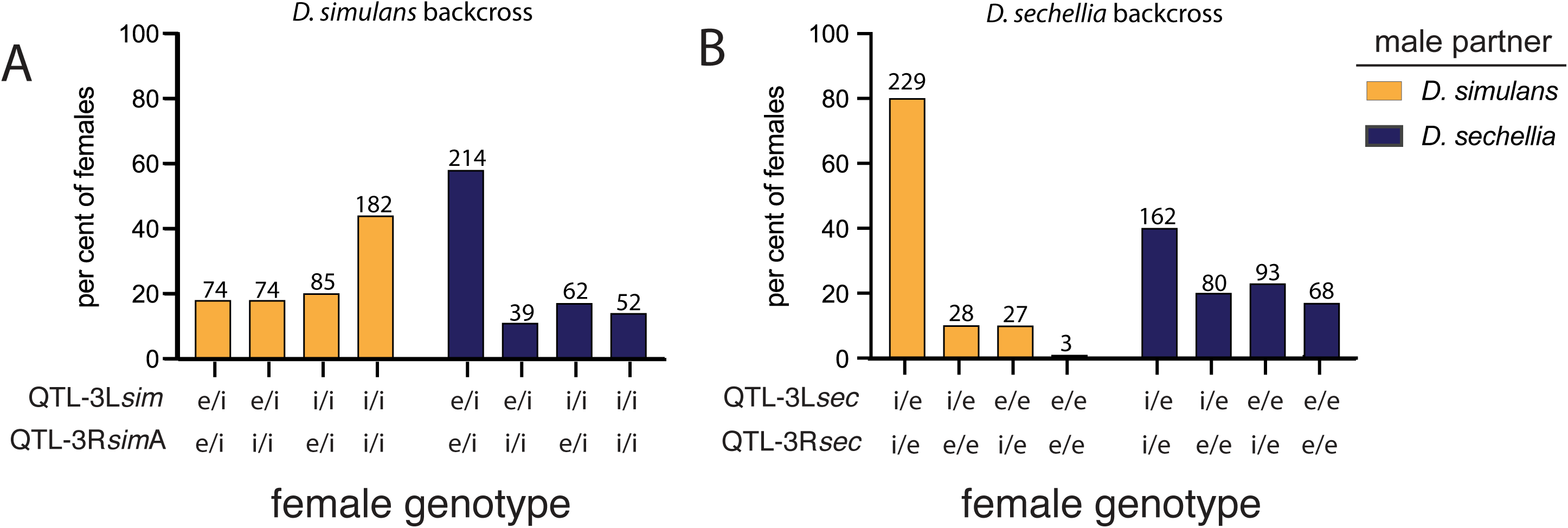
The percentage of females of each genotype at the two major QTL in the A) *D. simulans* backcross mapping, or B) *D. sechellia* backcross mapping population that had mated with *D. simulans* males (tangerine bars) or *D. sechellia* males (purple bars). The number of females of each genotype category is given atop each column. The genotype at 3L:11311968 was used as a marker of QTL-3L*sim* and the sequence at 3R:14758494 was taken for to genotype QTL-3R*sim*A, which are at the peak values. Similarly, we used the genotype at 3L:10480434 as the marker for QTL-3L*sec*, and the genotype at 3R:16634292as a marker for QTL-3R*sec*. e = *D. sechellia* allele and, i = *D. simulans* allele of each QTL.

Composite interval mapping (Figure 3A, grey lines) supports the same two major QTL peaks on 3L and 3R, but does not indicate additional significant QTL. We next explored the fit of multiple QTL models for the *D. sechellia* backcross population and found that the best-fit model supports a locus at ∼ 3L: 10.48 Mbp, near the peak detected in our one-dimensional scan (QTL-3L*_sec_*) and a second at ∼3R:16.63 (QTL-3R*_sech_*; Figure 3A; Table 1).

As with the QTL from the *D. simulans* backcross, the effects of each of the major loci were significant. The substitution of a *D. simulans* allele at QTL-3L*_sech_* had a 36.5% effect upon the likelihood of mating with *D. simulans* (Figure 3B), and the effects of substitution at QTL-3R*_sech_* had an effect of 32.8% (Figure 3C). The similarity between the QTL map for mate choice in the *D. simulans* backcross and the *D. sechellia* backcross could suggest that alleles of the same genes influence mate choice in backcross populations. The QTL map of chromosome 3L and 3R for the *D. simulans* backcross and *D. sechellia* backcross overlap significantly (Figure S1 A, B). While there is overlap at the major QTL on 3R, there is additional complexity, perhaps due to the presence of an additional locus at the distal end that is only significant in the *D. simulans* backcross (Figure 2A Figure S1).

We detected considerable epistasis between QTL-3L*_sech_* and QTL-3R*_sec_*, among the female *D. sechellia* backcross progeny that had mated with *D. simulans* males (Figure 4B). We found that substitution of a single *D. simulans* allele at either QTL-3L*_sech_* or QTL-3R*_sech_* could increase the likelihood of mating with *D. simulans* males 9-fold, but females bearing *D. simulans* alleles at both QTL were 76X more likely to mate with *D. simulans* males than females that were homozygous for the *D. sechellia* alleles at both (Figure 4B). We rejected the hypothesis that this pattern could be explained by an additive model of QTL effects (*p* = 10^-34^). As was the case for females of the *D. simulans* backcross, very strong epistasis between the major QTLs is observed in those females mating against the direction of the backcross.

### Tests of major QTL effects in no-choice assortative mating tests

The effects of each QTL on mating rate, competition, or differences in the propensity to mate could be masked in a two-choice test, so we also examined allele frequency of the major QTLs on 3L and 3R in samples of backcross females that were paired with males from only one species. This would discern whether a female genotype is attractive to males of one species, or whether it is disfavored by males of the other species. In contrast to the assortative mating tests for our mapping population when we only genotyped those that mated, we genotyped those females that failed to mate in this experiment. The genes *desaturaseF* (*desatF*) and *elongaseF (eloF)* are located within 1-2 Mbp of QTL-3L and QTL-3R, respectively, and we used polymorphisms in these genes to distinguish between *D. simulans* and *D. sechellia* alleles using PCR.

While mating occurred among all genotypes of the *D. simulans* backcross, crosses between female progeny of the *D. sechellia* backcross with males of either species are characterized by low mating rates among all genotype classes, and very strong repellence between some female genotypes and *D. simulans* males. Females that were homozygous for the *D. sechellia* alleles at both loci were the least likely to mate with either *D. simulans* or *D. sechellia*. No females that were *e/e; e/e* at QTL-3L*_sech_*; QTL-3R*_sech_* mated with *D. simulans* males, and strikingly, only about 9% of *e/e; e/e* females mated with *D. sechellia* males (Figure 5). Moreover, those females that were heterozygous at QTL-3L*_sech_* and QTL-3R*_sech_* (*i/e; i/e*) were the most likely to mate with *D. sechellia* males, even in the *D. sechellia* backcross (*p* = 0.005, two-tailed FET). These results confirm the patterns observed in the two-choice data but also reveal that for the females of the *D. sechellia* backcross, a lack of mating plays a larger role than competition for or between males.

**Figure 5.**
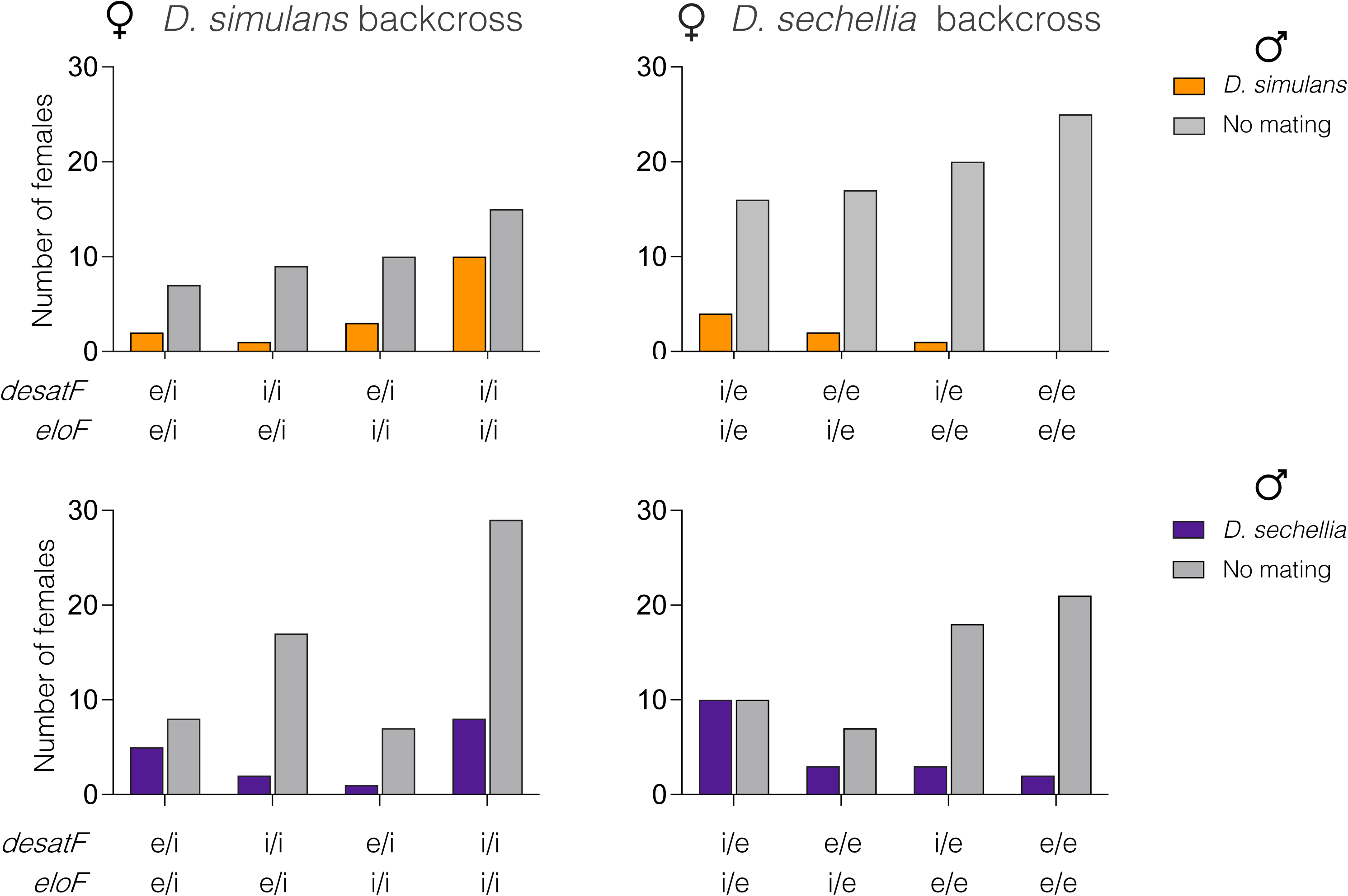
Genotypes at *desatF* (near QTL-3L) or *eloF* (near the major QTL on 3R) of females that successfully mated in no-choice tests with either *D. simulans* males (tangerine) or *D. sechellia* males (purple). The frequency of females with a given genotype that did not mate in the trials is shaded gray.

The assortative mating pattern for the no-choice tests of females of the *D. simulans* backcross also did not differ significantly from results of two-choice tests. Females that were homozygous for *D. simulans* alleles at both QTL-3L*_sim_* and QTL-3L*_sim_*A (*i/i; i/i*) were most likely to mate with *D. simulans* males. Females of all QTL-3L*_sim_* and QTL-3R*_sim_*A genotypes would mate with males of either species, but at different rates (Figure 5).

## Discussion

The sibling species *D. simulans* and *D. sechellia* have been an important model of speciation, and with the report of ongoing hybridization in the Seychelles (Matute and Ayroles 2014), they provide an opportunity to study species maintenance. A couple of pre-genome era studies have examined the genetic basis of reproductive isolation in females of these two species. Using insemination (Coyne 1992) or the production of offspring (Chu et al. 2013) the significant genetic contributions mapped to the X, 2^nd^ and 3^rd^ chromosomes.

We restricted our analysis to the behavioral components of isolation, with female choice and female attractiveness in the two-choice assay. In contrast to the previous studies that included insemination or production of offspring, we learned that behavioral isolation in females is localized to two major loci, one on each arm of the third chromosome. The QTL maps of assortative mating bear a strong resemblance to the maps of species-specific CHC production (Gleason et al. 2005; Gleason et al. 2009), suggesting that CHCs are the first, or primary behavioral barrier between *D. simulans* and *D. sechellia*.

Each of the two major-effect QTL for female assortative mating found in both backcross directions individually account for 32-37% of the difference in phenotype, which is exceptionally significant for a behavioral trait. The strength of the QTL occurs through two types of effects: 1) a difference in the relative mating rates with males of either species and, 2) whether the pairs mated at all. Females of all genotypes produced in the *D. simulans* backcross mate with males of both species, but *D. simulans* males are far more likely to mate with those females that are homozygous for the *D. simulans* alleles at QTL-3L*_sim_* and QTL-3R*_sim_*A (*i/i; i/i*) than with females bearing a *D. sechellia* alleles at either locus (Figure 4, 5). *D. sechellia* males, conversely, mate more readily with females bearing a *D. sechellia* allele at either QTL-3L*_sim_* or QTL-3R*_sim_*A but will mate with (*i/i; i/i)* females almost as readily (Figure 4A, 5).

Interactions in the *D. sechellia* backcross are more surprising. Females that are heterozygous at both QTL-3L*_sech_* (*DesatF*) and QTL-3R*_sech_* (*eloF*) were the most likely to mate with males of either species (Figure 5). Females that were homozygous (*e/e; e/e*) at both major QTL had the lowest mating frequency with either male. *D. sechellia* males will mate with *e/e; e/e* females only about 10% of the time in either no-choice or two-choice tests. On the other hand, *e/e; e/e* females never copulated with *D. simulans* males in the 25 no-choice tests (Figure 5) and only three of 691 (0.4%) copulations in the two-choice screen were between *e/e; e/e* females and *D. simulans* males (Figure 4). The strong barrier between *D. simulans* males and *e/e; e/e* females, therefore, underlies the significance of QTL-3L*_sech_* and QTL-3R*_sech_*.

### Parallels with the QTL map for 7,11-HD production

Our screen was designed to capture both the loci that contribute signals for male preference, and the loci that confer mate choice in females. However, the QTL map produced by our one-dimensional analysis of assortative mating in females bears a strong resemblance to a QTL map of variation in 7,11-HD production between *D. simulans* and *D. sechellia* females (Coyne et al. 1994b; Gleason et al. 2005; Gleason et al. 2009). For Gleason et al (2005; 2009) studies, two major QTL contribute to variation in 7,11-HD levels: one between markers near 3L:10.85 and 3L:11.88, and a second between markers near 3R:13.89 and 3R:15.42. These QTL co-localize with the major QTL that we have found, as well as with two enzyme-encoding genes that are required for conversion of 7-T to 7,11-HD, *eloF* (Chertemps et al. 2007; Combs et al. 2018) and *desatF (Chertemps et al. 2006; Shirangi et al. 2009)*. Genes for both enzymes are expressed in *D. sechellia* females but not in males, or in either sex of *D. simulans (Chertemps et al. 2007; Chertemps et al. 2006; Shirangi et al. 2009)*. 7,11-HD induces courtship of *D. simulans* females by *D. sechellia* males, but it inhibits courtship from *D. simulans* males (Billeter et al. 2009; Savarit et al. 1999).

The epistasis between our 3L and 3R major effect loci could be due to alleles of *desatF* and *eloF*, respectively, which together confer production of 7,11-HD in *D. sechellia* females. A similar epistatic effect between 3L and 3R loci has been described for production of 7,11-HD in *D. simulans* backcross progeny (Gleason et al. 2009); appreciable 7,11-HD levels are made when both *D. sechellia* alleles of *desatF* and *eloF* are present (Chertemps et al. 2007; Chertemps et al. 2006; Combs et al. 2018; Pardy et al. 2019). For *D. melanogaster* males, which are also attracted by 7,11-HD, the intermediate biochemical products that are produced when only one of the two enzymes are produced have no effect upon the attractiveness of females (Chertemps et al. 2007; Chertemps et al. 2006; Combs et al. 2018). Similarly, intermediate products produced by *D. sechellia* or *D. melanogaster* females that express either *desatF* or *eloF*, but not both, do not provoke aversion by *D. simulans* males (Combs et al. 2018). Therefore, acquisition of a single alleles of either *desatF* or *eloF* from *D. sechellia* will not cause a difference in 7,11-HD levels or a subsequent shift in female attractiveness on their own.

Is the barrier between *D. simulans* males and *D. sechellia* females due entirely to male *D. simulans’* aversion to 7,11-HD produced by *D. sechellia* females? We do not believe so. First, crosses between *D. simulans* and *D. melanogaster* are more successful when male *D. simulans* is paired with female *D. melanogaster*, which also produce 7,11-HD, but the reciprocal cross is rarely successful (Carracedo et al. 1998). If *D. simulans* males are entirely thwarted by 7,11-HD, they would be similarly thwarted in crosses with *D. melanogaster* females. Secondly, 7,11-HD is detected by contact chemoreceptors, not through olfactory signals (Seeholzer et al. 2018; Thistle et al. 2012). However, male *D. simulans* shun *D. sechellia* females in courtship assays, even without any physical contact between male and female (D. E., *manuscript in preparation*). Finally, many studies have shown that female Drosophila perceive male courtship song, and some have found that species-specific songs matter. For instance, *D. sechellia* females will mate with *D. simulans* males in 96-hour tests, but only if their wings have been removed (Tomaru et al. 2004; Ritchie et al. 1999). Therefore, sex– and species-specific difference in contact pheromones are unable to explain every feature of this behavioral boundary. So, if the QTL that we have uncovered here at peaks on 3L and 3R do correspond to alleles of genes that confer 7,11-HD production, it could mean that contact pheromones are either the strongest or the first in a series of species-specific checkpoints.

One candidate locus conferring female preference has also been mapped to the third chromosome in *D. simulans*. The *fruitless* (*fru*) gene is a ‘master-regulator’ of male sexual behavior in *D. melanogaster*. Deletions in the *fru* gene that unmask the *D. simulans* allele in *D. simulans/D. melanogaster* F1 hybrid females reveal that this locus confers female preference for *D. simulans* males and rejection of *D. melanogaster* males (Chowdhury et al. 2020; Laturney and Moehring 2012). Our one-dimensional maps do show a minor rise in LOD score around the *fru* locus (∼7Mbp, Sup. Figure 1). However, this region is not significant in either composite interval mapping or in multiple qtl models (Supp. Figure 1; Table 1), suggesting that in *D. simulans* females, *fru* imparts rejection of *D. melanogaster* males through a mechanism that is distinct from rejection of *D. sechellia* males.

### A “crowded” third chromosome

Loci that confer aspects of isolation between species are often found clustered together in the genome (Byers et al. 2021). The Drosophila genome is distributed over three major chromosomes (X, 2 and 3) and a diminutive ‘dot chromosome’, the fourth. Although the third chromosome is the largest, ∼36% of the genome, it still harbors a disproportionate number of loci that confer adaptive differences and reproductive isolation (Figure 6). A large body of work over decades has localized male mate choice to the right arm of chromosome 3 in a region near the gene *doublesex*: 1) a major effect locus for male pheromone (Coyne 1996), male courtship song (Gleason and Ritchie 2004), male mating success and courtship latency (Cantor and Civetta 2003) and courtship vigor (Shahandeh and Turner 2020). The region has been fine-mapped by Shahandeh and Turner (2020), who first remarked that the third chromosome is “crowded” because a cluster of three loci that together account for ∼44% of the variation in species-specific male ‘courtship vigor’ is located in and among QTL that confer 7,11-HD production in females (Shahandeh and Turner 2020). We showed that variation in female assortative mating, which should include female preference as well as female attractiveness, maps entirely to the third chromosome, and the entire chromosome is significant in our analyses.

**Figure 6.**
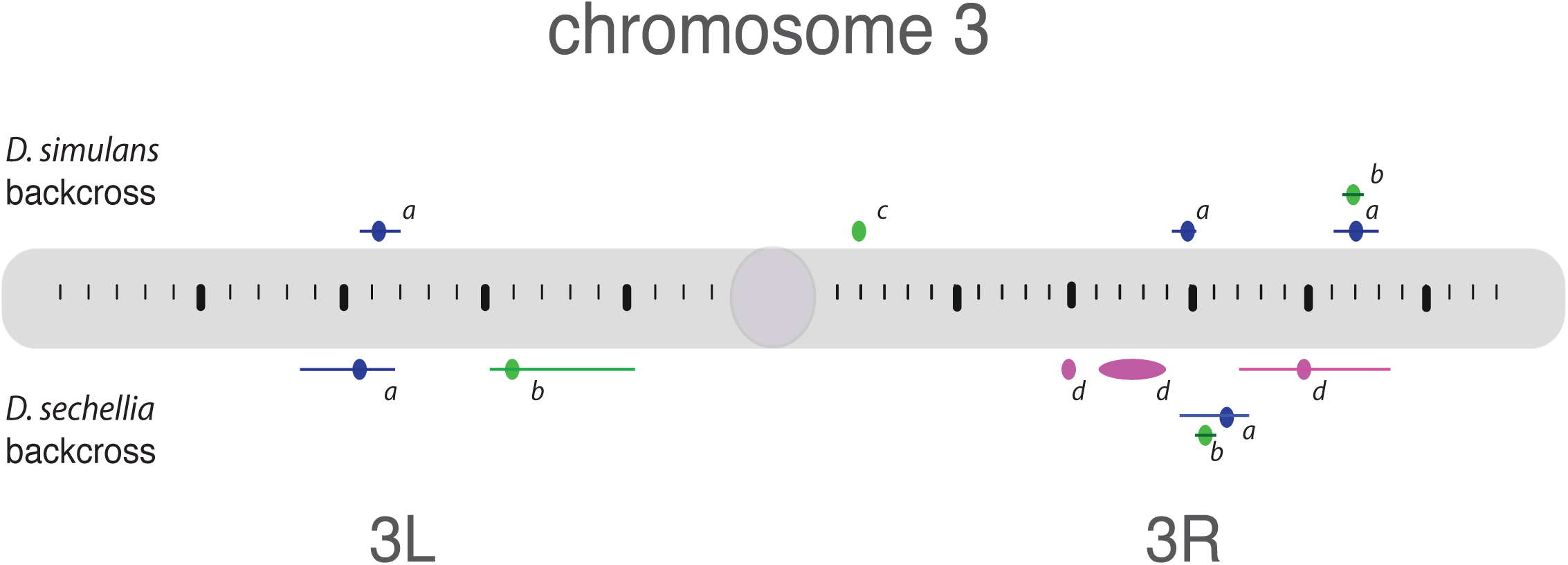
Loci that affect assortative mating in females are found inter-digitated among loci that confer resistance to the *M. citrifolia* fruit toxin, and with loci that confer species-specific mate preference in males. We include loci that confer either attraction to *M. citrifolia* or resistance to its toxin among the set of adaptive loci to those that 1) have been identified by mapping variation between *D. simulans* and *D. sechellia*, 2) have been localized to a few Mbp in the genome (rather than chromosome arm), and 3) confer at least 25% difference in the phenotype, and so are considered major-effect(Bradshaw et al. 1998). Bars represent 1.5 LOD confidence intervals. **a)** locations (in blue) of QTL-3L*_sim_* (∼ 3L:11.31 Mbp), QTL-3R*_sim_*A (∼3R:14.76) Mbp, QTL-3R*_sim_*B (∼3R:22.15), QTL-3L*_sech_* (∼3L: 10.48 Mbp) and QTL-3R*_sech_* (∼3R: 16.63 Mbp), this study; (**b**) the major effect loci conferring larval resistance (in green) in the *D. simulans* backcross (near the *Twdl* locus ∼3R:22.08 Mbp), and at ∼3L: 15.7Mbp and ∼3R:15.1 in the *D. sechellia* backcross (Huang and Erezyilmaz 2015) (**c)** *Osiris* locus (in green) conferring tolerance to *M. citrifolia* fruit toxin to adults (Hungate et al. 2013; Earley and Jones 2011a) (**d**) species-specific male courtship vigor maps to three loci on the third chromosome (in purple): ∼ 3R: 9.2-10.1 Mbp and ∼ 3R:11-13.9 Mbp and ∼ 3R:20.0 Mbp (Shahandeh and Turner 2020)

Of the seven major loci that contribute to adaptation to *M. citrifolia* fruit that have been mapped, all but two are on the third chromosome (Figure 6). The female assortative mating loci, QTL-3R*_sech_*, QTL-3R*_sim_*A and QTL-3R*_sim_*B are in a ∼7.4 Mbp cluster with two major effect loci for *M. citrifolia* toxin resistance. QTL-3R*_sim_*B is just 70 kilobase pairs from the peak of a locus of major effect (> 44%) for larval resistance (Huang and Erezyilmaz 2015). The peak of QTL-3R*_sech_* is just ∼1.5 Mbp from another larval resistance locus of significant effect (Figure 6). The left arm houses the female assortative mating loci QTL-3L*_sim_* and QTL-3L*_sech_* and these are within ∼5 Mbp of a major effect resistance locus (effect size = ∼40%) in larval resistance to *M. citrifolia* fruit toxin (Huang and Erezyilmaz 2015). In sum there are two clusters of mate choice and resistance loci. One is on 3R in a ∼12 Mbp stretch that also includes three loci conferring male courtship intensity, and another loosely linked association on the left arm of chromosome three. Another major effect locus conferring *M. citrifolia* resistance to adults is found near the centromere on 3R (Earley and Jones 2011b). Most of the loci that isolate the two species, therefore, are found on the third chromosome, and the proximity of alleles conferring assortative mating in males and females with loci conferring *M. citrifolia* resistance, could have been a factor in the initial divergence between species as well as maintenance of species boundaries during ongoing hybridization between wild *D. simulans* and *D. sechellia* in the Seychelles.

### Mating asymmetry and hybridization

*D. sechellia* can reproduce on fruit other than *M. citrifolia* and hybridization is currently underway in the Seychelles (Matute and Ayroles 2014). Hybridization between incipient species after secondary contact may reinforce isolation (Coyne and Orr 1989) promote range expansion, (Pfennig 2007)create ‘hybrid swarms’ (Moran et al. 2021), cause ‘despeciation’ (Turner 2002) or even produce new species (Lamichhaney et al. 2018). We find that the genetic architecture that underlies assortative mating in females of *D. simulans-D. sechellia* also promotes hybridization. In each of our experiments we found that females that mated with *D. simulans* males had more *D. simulans* alleles at QTL-3L and QTL-3R, but the reciprocal was not true for females that had mated with *D. sechellia* males. We found that *D. sechellia* males were more likely to mate with females that were heterozygous QTL-3L and QTL-3R in both the two-choice and one-choice datasets. Specifically, female progeny from the *D. sechellia* backcross that bear *D. simulans* alleles at both QTL-3L*_sech_* and QTL-3R*_sech_* (*i/e*; *i/e*) mated with *D. sechellia* males twice as often as with *e/e; e/e* in a two-choice assay. In the no-choice experiments the effect was more pronounced: those females that bore *D. simulans* alleles at one or both QTL-3L*_sech_* and QTL-3R*_sech_* (*i/e*; *i/e)* were more likely to mate when paired with *D. sechellia* than those females that were homozygous at both alleles (*e/e; e/e*; FET *p* = 0.005). Hence, the genetic architecture of assortative mating in females bears features that both promote integrity of the third chromosome of *D. sechellia* and features that promote mating with hybrids. Future field work on *D. sechellia* populations in the Seychelles will help to predict the fate of this endemic species.

## Data Availability

All strains of Drosophila are available upon request. Raw sequence reads for the two reference genomes and the separate genotyped library of hybrids will be deposited before publication on NCBI’s sequence read archive (SRA), indexed with a dedicated BioProject number. The genotype x phenotype files are available at: https://doi.org/10.5285/8dde3529-1cf7-4b0f-907d-a1631f38afd7 and the raw data for mating rates are provided at: https://doi.org/10.5285/361621ad-6487-47a4-bf8b-00f78705e593

## Acknowledgements.

We thank Christopher Black and Tausif Hasan of NYGenomes, and Thomas Scot Collins formerly of NY Genomes, who helped to install the MSG software for initial analyses. We are grateful to David Stern (Janelia, HHMI) for running the final analyses on updated MSG software. We also thank Prof. Chris Herzog (Kings College, London, Department of Mathematics) for calculating the significance of epistasis. Yan Huang, Karishma Suchday, Christie Yie, Emily Mann, and David Liu helped to produce and phenotype the recombinant females. Mike Ritchie (U. St Andrews) kindly provided helpful comments on an early version of the manuscript and members of the Goodwin Lab provided thoughtful discussion. This work was supported by Stony Brook University startup funds, as well as Stony Brook URECA fellowships to Christie Yie and Kenneth Lu. Deniz Erezyilmaz was supported by an AAUW Postdoctoral Leave Fellowship, and NERC Grant NE/S010351/1 to S.F. Goodwin.

## Supplementary Online Material

**Figure S1**. Overlay of LOD scores x physical distance (Mbp) of the third chromosome using data produced by **A**. Interval Mapping and **B**. Composite Interval. Mapping of the *D. simulans* backcross (solid lines) and the *D. sechellia* backcross (dashed lines). The *p* = 0.05 significance threshold is LOD 2.52 for the *D. simulans* backcross, and p = 2.57 for the *D. sechellia* backcross.

**File S1.** Calculation of epistasis.

**Supplementary Figure 1.**
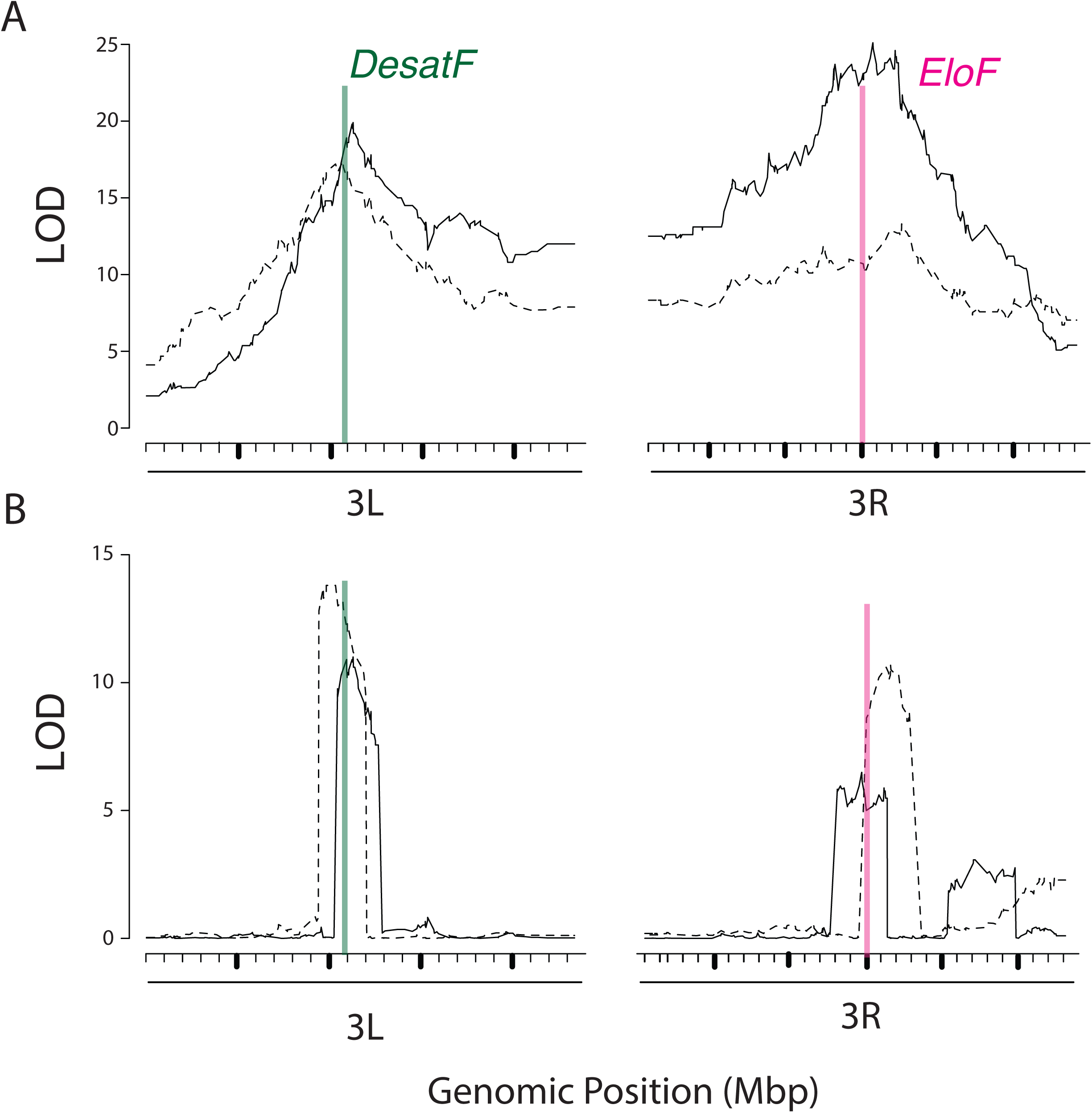
Overlay of LOD scores x physical distance (Mbp) of the third chromosome using data produced by A. Interval Mapping and B. Composite Interval. Mapping of the *D. simulans* backcross (solid lines) and the *D. sechellia* backcross (dashed lines). The *p* = 0.05 significance threshold is LOD 2.52 for the *D. simulans* backcross, and *p* = 2.57 for the *D. sechellia* backcross. The locations of *DesaturaseF(DesatF)* and *ElongaseF (EloF*) are marked in green and pink, respectively.

We consider a system of two genes, each of which conjecturally affects a particular phenotype, and where the organism is in one of the following four states:

(A a) (B b)

(A A) (B b)

(A a) (B B)

(A A) (B B)

The null hypothesis is that these two genes do not interact with each other. If the phenotype can be measured by a number E, then this hypothesis leads to the following quantitative statement about E. There is a baseline effect E_0 that contributes to the phenotype regardless of genotype. The (A A) genotype will lead to an increase in the effect by E_A (or a decrease if E_A is negative). The (B,B) genotype will lead to an increase in the effect by E_B. Thus, we guess that in the four cases, we should have an effect of the following form

(A a) (B b): E_1 = E_0

(A A) (B b): E_2 = E_0 + E_A (A a) (B B): E_3 = E_0 + E_B

(A A) (B B): E_4 = E_0 + E_A + E_B

This model leads to the following prediction on the linear combination

E_1 + E_4 – (E_2 + E_3) = 0

Now, how do we test if the model is right? For each of the measured effects, E_1 through

E_4, we have an associated error \delta E_1 through \delta E_4. The errors are correlated and need to be summed carefully:

\delta E = \sqrt{ 4 \delta E_2^2 + 4 \delta E_3^2 }

We are using here that E_1 + E_2 + E_3 + E_4 = sample size and so E_1 + E_4 – (E_2 + E_3) = sample size – 2 E_3 – 2 E_4

with presumably negligible error on the sample size. The data will give some result

R = E_1 + E_4 – (E_2 + E_3)

which is presumably not exactly zero. By dividing R by \delta E, we find out how many standard deviations we are away from zero. This number of standard deviations can be converted to a p-value in a standard way.

We estimate the errors on E_1 through E_4 by assuming you have a normal distribution of flies (i.e. your error just comes from statistical sampling error from taking flies from this distribution).

There is a standard formula for the individual errors

\delta E_1 = \sqrt{ \delta E_1 (1 – \delta E_1) / sample size} assuming \delta E_1 is expressed as a number between 0 and 1.

I have found the following results for your data sets

set 1:

sample size = 287;

effect size in percent = {79.79, 9.76, 9.41, 1.05};

\delta E_i in percent = {2.4, 1.8, 1.7, 0.6}

\delta E in percent = 4.9 R = 61.3

p = 3 x 10^{-34}

set 2:

sample size = 403;

effect size in percent = {40.2, 19.9, 23.1, 16.9}

\delta E_i in percent = {2.4, 2.0, 2.1, 1.9}

\delta E in percent = 5.5 R = 14.1

p = 0.056

set 3:

sample size = 415

effect size in percent = {17.83, 17.83, 23.1, 16.9}

\delta E_i in percent = {1.9, 1.9, 2.0. 2.4}

\delta E in percent = 5.5 R = 23.3

p = 0.0019

set 4:

sample size = 367

effect size in percent = {58.3, 10.63, 16.89, 14.17}

\delta E_i in percent = {2.6, 1.6, 2.0, 1.8}

\delta E in percent = 5.1 R = 45.0

p = 3 x 10^{-17}

The conclusion is that sets 1 and 4 are much more poorly modeled by this additivity assumption.

footnote:

The formula I used to calculate p was

p = 2 \int_{R / \delta E / \sqrt{2}}^\infty x^2 \exp(–x^2) dx / \sqrt{\pi} (two tailed p value)

